# Relatedness modulates reproductive competition among queens in ant societies with multiple queens

**DOI:** 10.1101/2021.12.19.473293

**Authors:** Heikki Helanterä, Martina Ozan, Liselotte Sundström

**Affiliations:** Faculty of Biological and Environmental Sciences, Organismal and Evolutionary Biology, P.O.BOX 65, Helsinki University, FI00014 Finland; Tvärminne Zoological station, J.A. Palménintie 260, 10900 Hanko, Finland; Faculty of Science, Ecology and Genetics research unit, University of Oulu, FI 90014 Finland

**Keywords:** *Formica fusca*, reproductive skew, kin discrimination, inclusive fitness, conflict, cooperation

## Abstract

Reproductive sharing in animal groups with multiple breeders, insects and vertebrates alike, contains elements of both conflict and cooperation, and depends on both relatedness between co-breeders, as well as their internal and external conditions. We studied how queens of the ant Formica fusca adjust their reproductive efforts in response to experimental manipulations of the kin competition regime in their nest, as well as their own reproductive status. Queens respond to the presence of competitors by increasing their egg laying efforts, but only if the competitors are highly fecund and distantly related. Furthermore, queens only engage in cannibalism of eggs when the risk of erroneously destroying own offspring is absent. We demonstrate that queens of Formica fusca fine-tune their behaviours in response to kinship and fecundity of others as well as their own physiological state in an unusually precise manner.

## Introduction

Reproductive cooperation often involves individuals with diverging fitness interests, so that reproductive conflicts among them arise. A multitude of ecological, genetic, and social factors have been proposed to shape the outcome of reproductive conflicts and the relative reproductive success of cooperatively breeding individuals (i.e. reproductive skew; Reeve and Keller 2001). The strategies used for increasing reproductive shares depend on the kinship between group members, differences in their reproductive traits, and the availability of information about these. For example, in dwarf mongooses, *Helogale parvula*, the dominant female inhibits ovulation of subordinates via aggression (Creel et al. 1992), by which she is able to almost entirely monopolize reproduction in the group. Conversely, competitive reciprocal destruction of each other’s eggs by females of the cooperatively breeding acorn woodpecker, *Melanerpes formicivorus* equalizes reproductive shares (Koenig et al. 1995), believed to promote synchrony in egg laying and reduce reproductive skew. Such interactions may be modulated by relatedness in complex ways, such as when dominant banded mongoose females preferentially harass and even evict closely rather than distant kin, as close kin are less likely to aggressively retaliate (Thompson et al 2017).

In eusocial insects, reproductive division of labour between morphologically distinct castes permits direct fitness to only a minority of individuals i.e. the queen(s), whereas the majority of individuals, i.e. the non-reproductive workers, largely only gain indirect fitness returns (Hamilton 1964). Colonies of many ant species furthermore contain several reproductively active queens (polygyny). Reproductive conflicts may arise among these owing to limited colony resources (Ross 1988; Fournier et al. 2004; Bargum and Sundström 2007a; Kümmerli and Keller 2007). The reproductive shares of queens may be unequal (Hannonen and Sundström 2002; Fournier et al. 2004), but this has usually been established only at the pupal stage (but see Ross 1988). Thus, the question at what stage such differences arise, and how, remains uncharted.

The reproductive shares of queens may be affected by several factors. First, queens may differ in intrinsic traits, such as age, egg laying rate, and egg viability (Ross 1988, Hughes and Strassmann 1988; Hannonen and Sundström 2002; Holzer et al. 2006). The shares may also reflect temporal variation in fecundity, whereby queens that start oviposition early gain larger shares of reproduction to the detriment of late starters (Ozan et al. 2013). Second, queens may compete over reproduction via overt aggression or brood cannibalization. The former may be rare in secondarily polygynous ant colonies, i.e. those where multiple queens co-exist in mature colonies (Keller 1993), but the latter has been observed in e.g. *Ectatomma* (Hora et al. 2007), and *Leptothorax* (Bourke 1994). Cannibalism may also be moderated by differences in the onset of ovipositioning among individuals, so that by cannibalizing eggs before the onset of own oviposition, a queen will minimize the risk of eating own progeny (Hora et al. 2007). Third, queens may chemically suppress oviposition of other queens, as found in both ants (Vargo 1992, Holman et al 2013, Abril & Gomez 2020), and termites (Yamamoto and Matsuura 2011). Finally, indirect genetic benefits mediated by relatedness patterns may influence interactions among nest mate queens, (Hamilton 1964). In secondarily polygyne ant colonies the relatedness among queens can vary extensively (Stuart et al. 1993; Brown and Keller 2002; Hannonen and Sundström 2003; Kikuchi et al. 2007; Vásquez and Silverman 2007; Meunier et al. 2011). This may create an incentive for queens to adjust their responses to competing queens and so balance direct fitness gains from own reproduction with indirect fitness gains via related queens.

In this study we carry out experiments to tease apart the different factors that contribute to reproductive sharing among nestmate queens in the ant *Formica fusca*. Colonies of this species often have several queens, with variable degrees of relatedness (Hannonen & Sundström 2003). The queens do not compete by overt aggression, yet differ considerably in both fecundity and their ultimate reproductive shares (Hannonen and Sundström 2002; Ozan et al. 2013). In the first experiment, we investigate whether queen fecundity changes in response to the presence of odours of another queen, and whether the response depends on fecundity and relatedness of the other queen, as would be expected if the queens communicate or suppress each other chemically. In the second experiment we investigate whether queens cannibalize eggs of other nest mate queens and whether this varies according to queen reproductive status and relatedness to brood. We expect the queens should cannibalize eggs at a higher rate if they have themselves not yet reproduced and/or the relatedness to brood is low.

## Methods

### Colony collection and maintenance

We collected 30 polygyne colonies of the black ant, *Formica fusca*, on the Hanko peninsula, south west Finland in late April and early May 2012. The colonies were sorted to determine queen number, and to ensure that oviposition had not yet commenced, after which they were stored at +4°C with nest material and peat for 1-2 weeks, until the experiments started. These conditions emulate continued hibernation and prevent queens from resuming reproduction before the onset of experiments. Once enough colonies were available for the experiments, the ants were transferred to ambient temperature (22-24°C) to prepare them for the experiments. At this stage, the nests were housed in plastic trays (30×40×15cm) with plaster bottom, and the walls coated with Fluon™. Water was provided in Eppendorf tubes plugged with cotton, and the ants were fed standard Bhatkar diet (Bhatkar and Whitcomb 1970) daily.

#### Experiment 1 – Competition via chemical communication

To investigate whether queen fecundity changes in response to chemical cues that signal the presence of other queens, we selected colonies with three or more queens, and divided them into experimental nests with single queens, each with an entourage of ca 50 workers (Figure S1). After five days we counted the number of eggs each queen had laid, to assess their fecundity. We then randomly selected one queen (donor queen), which we killed by freezing and extracted the non-volatile cuticular compounds, to be used as chemical cues for the presence of another queen in the colony. The whole-body surface chemicals of the donor queen were extracted in 200µl Chromasolv pentane (Sigma-Aldrich). The first extraction volume was allowed to evaporate and the dissolved chemicals re-diluted in 160µl pentane. After extracting the cuticular compounds, the donor queens were placed in 94% ethanol for genotyping.

The two remaining queens were placed individually on Petri dishes without workers overnight before the experiment started (Figure S1). If the queens laid eggs during this period, we added those to the egg counts determined during the first five days of worker presence. The next day the queens were moved to plastic jars (Ø 7cm), lined with plaster, and the walls coated with Fluon™. A cavity (Ø 3cm) covered with a piece of transparent red plastic sheet (2×5cm) served as a brood chamber. A glass oval-shaped bead (8×5mm) treated with either 10µl of extract from the donor queen (treatment queens) or pentane (control queens), was placed in the brood cavity to simulate the presence of another queen (Figure S1). The glass beads were treated every 12 hours for the duration of the experiment (five days). The queens received water, as described above, but no food during the experiment. At the end of the experiment, we counted the eggs the queens had laid, to determine queen fecundity during the experiment, and placed the treatment and control queens in 94% ethanol for genotyping.

To assess whether queen relatedness influences changes in queen fecundity we genotyped all donor, treatment and control queens. We used twelve polymorphic microsatellite loci designed for *Formica* species: FL12, FL20, FL21(Chapuisat 1996); FE13, FE16, FE19, FE21, FE42, and FE51 (Gyllenstrand et al. 2002) and FY4, FY7, FY13 (Hasegawa and Imai 2004). Details of the protocols are given in the Supplementary information.

### Statistical analyses

We estimated the pairwise relatedness between treatment and donor queens based on the maximum likelihood estimation in the software ML-Relate (Kalinowski et al. 2006). To investigate whether the presence of a nest mate queen influences queen fecundity, we first tested the difference in egg-laying rate between the treatment queen and the control queen in each colony in the experimental phase using paired t-test. We then tested whether the fecundity of the donor queen, and her relatedness to the treatment queen induced changes in the fecundity of the treatment queen using a linear model (LM) in R (R Development Core Team 2014). Our continuous predictors were the fecundity of the donor queen and her relatedness to the treatment queen. Because the response to nest mate fecundity may depend on relatedness between the queens, we included the interaction between donor queen fecundity and relatedness in the model. Our response variable was the difference between the daily egg-laying rates in the experimental phase between the treatment and control queens.

#### Experiment 2 - Egg cannibalism

In this experiment we investigated whether queens selectively cannibalize eggs of nest mate queens, and whether queen reproductive status (non-reproductive vs. reproductive), and egg provenance (own or nest mate) affects this behaviour. To achieve this, we presented the queens with eggs of different provenance, and examined the number of eggs eaten or destroyed. We first selected 2-4 queens per colony, obtaining in total 71 queens from 30 colonies, using a different set of colonies than Experiment 1. One queen from each colony was maintained at +4°C in order to prevent egg laying (non-reproductive queen), whereas the remaining ones were placed individually at room temperature (22-24°C), and thus allowed to start laying eggs (reproductive queens). In both cases, each queen had an entourage of 20-25 workers. The setup of nests and conditions were identical to the initial stage of Experiment 1. Once enough eggs (min. 20-40) were present in the fragments with reproductive queens (4-5 days) to allow the set-up of eggs per colony as illustrated in Figure S2, the nest fragments with the non-reproductive queens were removed from cold storage, and the experiment started.

To ensure that queens could not replace introduced eggs with their own, we prevented all experimental queens from ovipositing during the experiment by covering their ovipositor with nail polish (IsaDora, Invima AB). Prior trials did not reveal adverse effects of nail polish on *F. fusca* queens. The nail polish could readily be removed from the cuticle by rubbing the queen ovipositor against a piece of moistened sponge. To apply the nail polish, we first immobilized the queen by chilling her in the freezer (−20°C) for ca. 4 minutes, and then pinned her down on a piece of polystyrene to allow easy access to the ovipositor. Once painted, we allowed 5 minutes for the nail polish to harden, during which time queen gradually regained activity. The queens were then placed individually in their respective experimental jars for 3-4 hours in the dark to calm down.

At the start of the experiment, we introduced 20 eggs into each experimental jar on the top of the sheet that covered the nest cavity (Figure S2) taking care to ensure that the eggs were intact. All non-reproductive queens were presented with eggs laid by one nest mate queen i.e. treatment A (Figure S2), whereas reproductive queens were presented either with eggs laid by a nest mate queen, an equal mixture of nestmate and own eggs randomly spread, or only their own eggs i.e. treatments B-D, respectively (Figure S2). Each experimental jar contained only one queen to be tested for cannibalism but several queens from the same colony were used in different treatments. The housing and conditions were identical to those in experiment 1, except that to maintain sufficient moisture, the jars were covered with a sponge, which was moistened daily. The experiment was terminated after five days, at which time the surviving queens (69%) were still active and had the nail polish intact. We counted the number of intact eggs that remained in the treatments with live queens and used the difference between the eggs introduced and those remaining as a measure of egg cannibalism.

### Statistical analysis

We compared queen survival in the four treatments (A-D) using a Chi-square test. We then compared egg survival across treatments using a Generalized linear mixed model (GLMM) with binomial errors in R (package lme4) (R Development Core Team 2014; Bates et al. 2015). The four experimental treatments were entered as categorical predictors, and the number of surviving vs. cannibalized eggs as the response variable. Because different queens from the same colony were used in different treatments, we added colony as a random variable in the analysis. We tested significances with the likelihood ratio-based χ^2^ test and compared mean egg survival among experimental treatments using Tukey contrasts (package multcomp; Hothorn et al. 2008).

## Results

### Experiment 1

The average daily egg-laying rate of all experimental queens before and during the experiment was 17.0±9.7 and 2.3±1.9 (mean±SD), respectively. The egg-laying rates of the treatment and the control queens during the experiment did not differ significantly from each other (paired t-test: t=1.51, df=14, *p*=0.153). This suggests that queen fecundity was unaffected by the presence versus absence of CHC odour of a nest mate queen. However, the egg-laying rate of the treatment queen (compared to that of the control queen) decreased with the fecundity of the donor queen, if queens were related. This relationship was reversed when the relatedness between the treatment and the donor queen was low (fecundity: t= −3.57, *p*=0.004; relatedness: t= −1.9, *p*=0.08; fecundity x relatedness: t=4.46, *p*=0.001; df=11, r^2^= 0.74; Figure 1).

**Figure 1.**
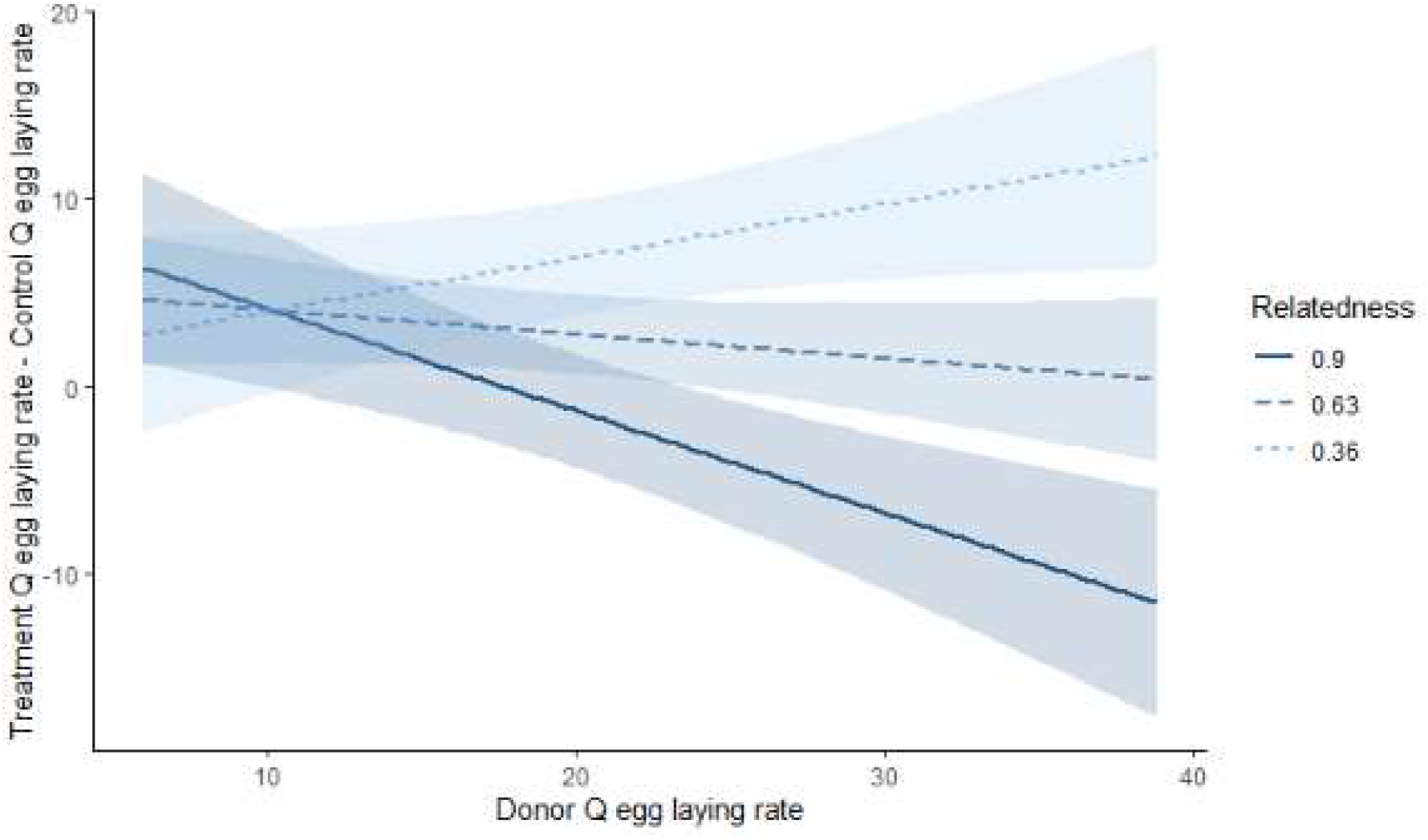
The effect of fecundity i.e. the egg-laying rate of (odour) donor nest mate on the fecundity difference between treatment and control queens as a function the relatedness between treatment and donor queens. The slopes (and 95% CI) of the relationship between predicted queen fecundity and the fecundity of her nest mate were estimated at high (mean+1SD), mean and low (mean-1SD) relatedness, using a simple slopes analysis (Dawson, 2014), using R-package *jtools* (Long et al. 2020).

### Experiment 2

In total, 49 queens from 24 colonies survived the experimental treatment. The survival of queens was lowest among non-reproductive queens (i.e. treatment A) (χ^2^= 8.30, df=3, *p*=0.040; queen survival by treatment: 8/18 queens (44%); 12/16 queens (75%); 15/17 queens (88%); and 14/20 queens (70%) in treatments A-D, respectively). However, the survival of reproductive queens between treatments (treatments B-D) did not differ significantly when non-reproductive queens were omitted from the analysis (χ^2^**=**1.82, df=2, *p*=0.403).

Egg cannibalism was very rare, as on average 19 out of 20 introduced eggs survived the experiment undamaged (19.0±1.9; mean±SD), yet egg survival differed between treatments (GLMM treatment χ^2^ =17.52, DF=3, *p>*0.001; Figure 2). Cannibalism by non-reproductive queens was significantly more common (treatment A), compared to all the other treatments (mean egg survival±SD by treatment: *A* = 17.3±3.9 (n=8 queens); *B* = 19.5±0.7 (n=12); *C*=19.6±0.5 (n=15) and *D*=18.9±1.6 (n=14) (contrasts *A vs. B-D* (min. and max. value): 3.62≥ *z* ≤2.66; 0.002≥ *p* ≤ 0.038) (Figure 2, Supplementary Table 1). Differences among treatments B, C and D were not significant (*z* ≤ 1.67, *p* ≥ 0.34).

**Figure 2.**
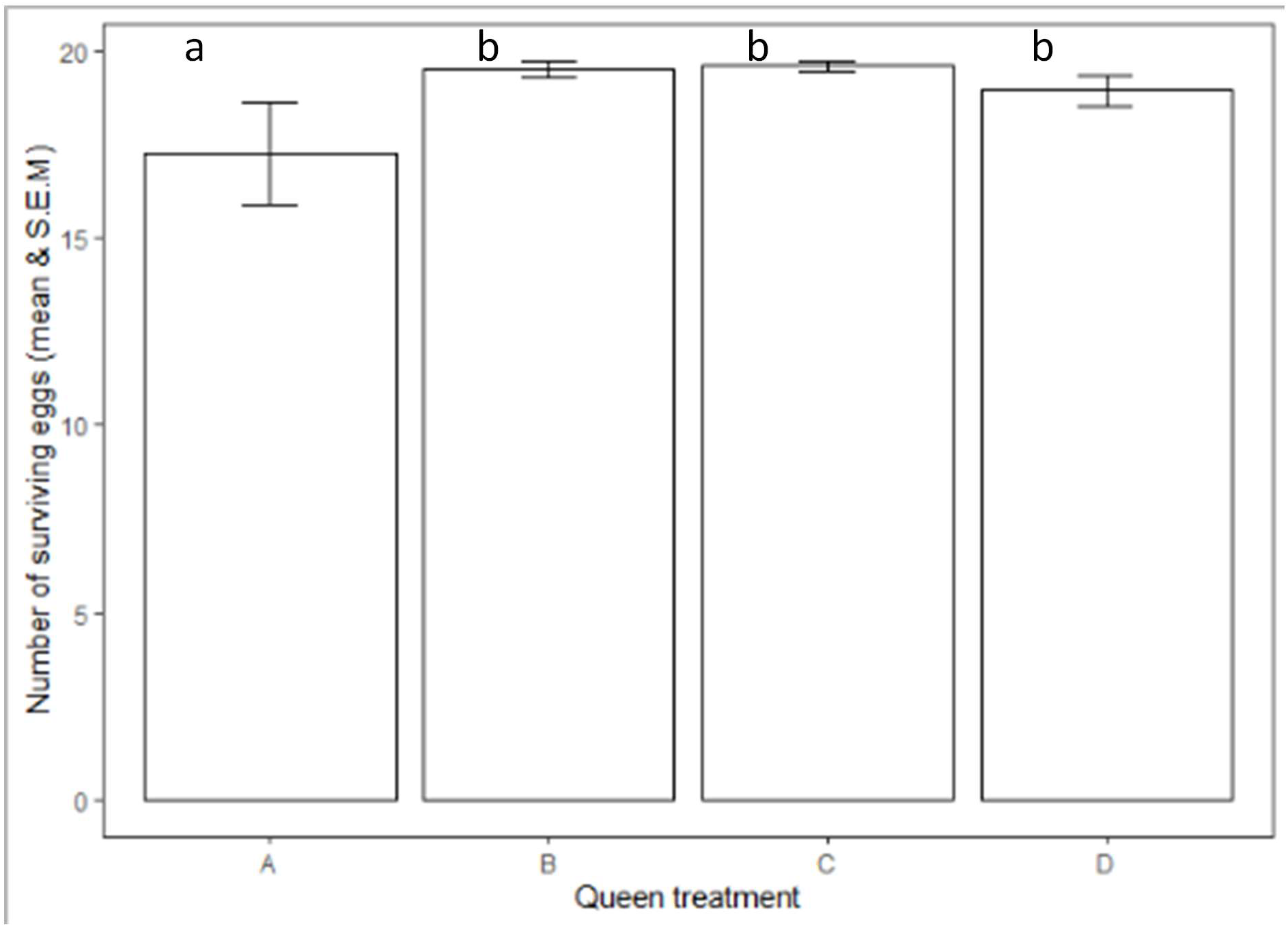
Egg survival at the end of experiment 2 (mean±95% CI). The letters A-D denote the treatment i.e. reproductive status of the experimental queen and the origin of the eggs offered. The letters “a” and “b” mark significant difference between treatments.

## Discussion

We tested whether queens adjust their egg-laying rate in response to chemical cues from competitors, and whether they cannibalize eggs to enhance their own reproductive output. We show that the response to competitor odours was modulated by fecundity and kinship. Highly fecund competitors had a negative effect on treatment queen fecundity when the queens were close relatives, but this effect was reversed when relatedness was low. This suggests that highly fecund queens are not able to simply coerce diminished egg laying in their competitors, but that queens respond to chemical signals from their nestmates in a facultative, kinship sensitive manner. Finally, in agreement with our predictions, we show that non-reproductive queens cannibalized more nest mate eggs than reproductive queens. This suggests that queens may suffer increased costs of cannibalizing once they themselves begin to reproduce.

We show a kinship sensitive effect of the odours of another queen. This is in contrast with observations from other social insects where the presence of a competing queen or her odours simply decreases fecundity (Vargo 1992; Yamamoto and Matsuura 2011; Holman et al. 2010a, Abril & Gomez 2020 but see Bourke 1993; 1995), and queens in polygyne ant colonies have reduced per capita fecundity compared to those in single-queen colonies of the same species (van der Meer et al. 1992; Trunzer et al. 1998; Schrempf et al. 2011; Yamamoto and Matsuura 2011). The more complex facultative and kinship dependent response observed here in *F. fusca* could reflect the highly variable kin structures of the species (Hannonen et al 2004, Bargum et al 2007b, Ozan et al 2013), that create conditions where such adjustments potentially lead to inclusive fitness gains. Our results show similarities to those found in allodapine bee *Exoneura robusta*, in which queens inhibited their ovary development more in when in associations with related queens than with unrelated queens (Harradine et al. 2012). Such findings suggest that the chemical signals between queens may be mutually beneficial rather than coercive in nature, as in the honest signals that mediate effects of a queen on worker fertility (Holman et al 2010b).

Our results suggest that *F. fusca* queens respond facultatively to kinship associated odour cues in order to enhance their indirect fitness returns. While the odours on queen cuticle have not been analysed in *F. fusca*, eggs laid by queens have been shown to carry matriline-specific chemical cues in this and other *Formica* species (Helanterä & d’Ettorre 2015). Furthermore, workers are able to recognize their mother even when reared in foster nests (El-Showk et al. 2010), and to nepotistically favour closely related queens (Hannonen & Sundström 2003). Together these findings suggest that queens could be able to use chemical information to detect both the fecundity and kinship of odours presented to them, even if such kin discrimination within nests is rare in insect societies (Boomsma & d’Ettorre 2014).

Our results raise the question why queens should trade-off direct reproduction for indirect in the presence of related queens. Possible reasons include avoidance of indirect fitness costs of competition with kin (West et al. 2002), and direct costs of sub-optimally high reproductive investment on future reproduction or survival. As such direct costs have not been observed in ants in recent studies (Heinze and Schrempf 2012; Heinze et al. 2013; von Wyschetzki et al. 2015), we suggest that reproductive allocation may be optimized at the colony, rather than individual level in the high relatedness setting. It is possible that the queens, for example, are able to lay more eggs later in the season to increase production of future workers in the nests. This might benefit all co-habiting queens, and pay off under high kinship, as has been observed in this species (Bargum et al. 2007ab).

Despite egg cannibalism being rare in our experiment, non-reproductive queens cannibalized more often than reproductive queens. This suggests that costs of indiscriminate egg cannibalism may lower for queens than have not yet began to oviposit, whereas reproductive queens may avoid egg cannibalism, given that they seem unable to discriminate between own and nestmate eggs. Egg destruction by non-reproductive individuals is a relatively wide spread phenomenon in joint nesting and cooperatively breeding birds (Koford et al. 1990; Koenig et al. 1995; McRae 1996; Macedo and Bianchi 1997) and communally breeding insects (Eggert 2000), and confers benefits without the necessity to recognize kin. Although some of the eggs introduced for cannibalism may have been laid by workers (Helanterä and Sundström 2005), this should not be confounded by treatment in our experimental design.

We show that despite their dependence on non-reproductive helpers for survival and brood care, queens can nevertheless mediate their reproductive interests by subtle means when reproduction is shared. *Formica fusca* queens are able to achieve this by cannibalizing the nestmate queen eggs before the onset of own reproduction and by responding facultatively to kin informative chemical cues on nest mate cuticle to increase inclusive fitness returns once queen begins to oviposit. Facultative reproductive adjustments based on relatedness have likewise been reported in cooperatively breeding vertebrates (Cornwallis et al 2009), but less so in social insects (but see Foster and Ratnieks 2000; Harradine et al. 2012; Ozan et al. 2013), and even more rarely among co-breeding queens. The ability of *Formica fusca* queens to actively shape their own reproductive effort before the onset of brood rearing suggests that queens in associations with apparent lack of overt reproductive conflicts have more power over their own reproductive destiny than may be generally appreciated, and that the balance between cooperation and conflict is fine-tuned to the kin structure context.

## Supporting information

DataFile2

DataFile1

DataFile3

## Notes

### Competing Interest Statement

The authors have declared no competing interest.

