## SupplementaryInformation for "Relatedness modulates reproductive competition among queens in ant societies with multiple queens"

**Supplementary information**

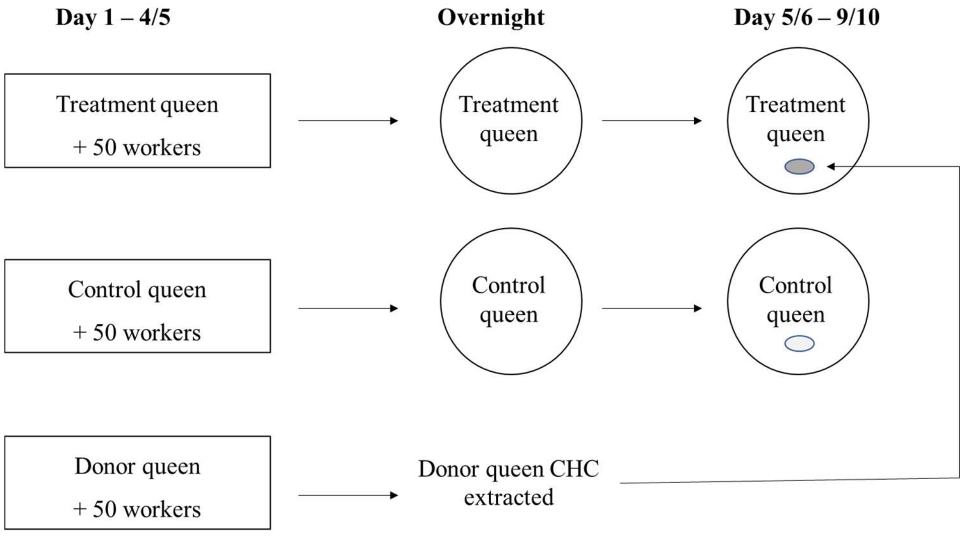


**Figure S1 –The experimental set-up for testing queen fecundity changes in the presence of a nest mate queen using a single three-queen colony. Following oviposition in the presence of workers (square boxes), the cuticular hydrocarbons of the donor queens were extracted. The two remaining queens were placed in isolation overnight, and then assigned to their experimental treatments (circles). The treatment queen was housed with a glass bead coated with the CHC of donor queen and control queen was housed with a glass bead treated with pentane control.**

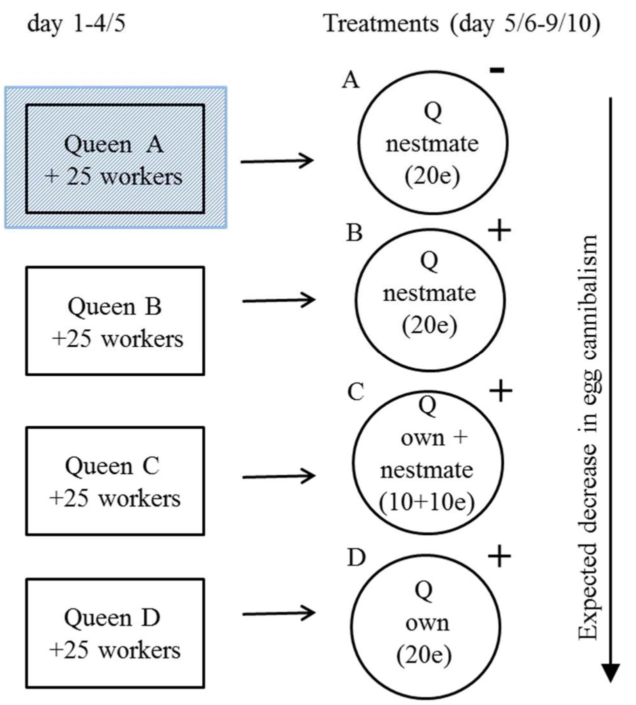


**Figure 2** – Experiment 2 **-** The experimental set-up for one colony to test the extent of egg cannibalism by nest mate queens. Square boxes indicate worker fragment containing reproductive queens before the experiment. The oviposition of queen A was inhibited by keeping her in cold (as indicated by pattern fill). Circles indicate experimental jars (i.e. treatments A-D) that differed in the composition of eggs (e) offered to queens to be cannibalized (Q). Queen reproductive status i.e. whether queen has oviposited or not prior to the experiment is denoted by +/- signs.

### Supplementary methods: DNA extraction and PCR methods

DNA was extracted by incubating two legs from each queen O/N at 56°C in 2.5:100µL proteinase K-Chelex (Schultner et al. 2014). PCRs were run in 10 µL reactions using 5 µL Qiagen Type-It microsatellite multiplex buffer, 3 µL deionized water, 1 µL optimized primer mix, and 1 µL DNA at 1:200 dilution on a 3730 ABI sequencer, following the protocols recommended by Qiagen. The microsatellite peaks were scored with Genemapper v. 4.1, and allele calling confirmed manually.

**Supplementary table 1 –** Differences in cannibalism levels between treatments.

Overall model: $\chi 2 = 17.52$; DF=3; $p>0.001$

| treatment contrasts | Estimate | Std. Error | $z$-value | $p$-value |
|---|---|---|---|---|
| A vs. B | 1.77 | 0.56 | 3.15 | 0.006 |
| A vs. C | 1.99 | 0.55 | 3.62 | 0.001 |
| A vs. D | 1.08 | 0.41 | 2.66 | 0.029 |
| B vs. C | 0.22 | 0.61 | 0.36 | 0.984 |
| B vs. D | -0.69 | 0.56 | -1.24 | 0.481 |
| C vs. D | -0.91 | 0.54 | -1.67 | 0.244 |

**Supplementary Information: description of data files**

**total_egglaying.txt**

Variables:
colony: ID of the original field collected colony where queens and workers originated from

queen: unique ID of the queen

role: whether the queen belongs to treatment or control

workers: present = queen was with 50 workers, absent = queen alone

egg-laying: eggs laid per day

**donor_eff.txt**

Variables:
col: ID of the original field collected colony where queens and workers originated from

q3pre: daily egg laying rate of the donor queen

Relatedness: relatedness between donor and treatment queen

q1post: egg laying rate of the treatment queen under exposure to odours from donor queen

q2post: egg laying rate of the control queen when exposed to pentane control

**canni.txt**

Variables:

Colony: ID of the original field collected colony where queens and workers originated from

Queen: treatment of the queen
A: non-reproductive queen with nestmate eggs
B: reproductive queen with nestmate eggs
C: reproductive queen with own and nestmate eggs
D: reproductive queen with own eggs

Alive: number of surviving eggs (out of 20) at the end of the observation

Dead: 20 - number of surviving eggs
