## Supplementary material for "Relatedness modulates reproductive competition among queens in ant societies with multiple queens": R code

library(multcomp)
library(ggplot2)
library(jtools)
library(plyr)
library(lme4)

total_eggs<-read.table("total_egglaying.txt", header=T)
# read data file with egg laying rates in the presence and absence of workers
attach(total_eggs)
tapply(egg.laying,workers,mean)
tapply(egg.laying,workers,sd)
# means and standard deviations of egg laying before and after the experiment

donor_effect<-read.table("donor_eff.txt", header=T)
 # read data file with egg laying rates in the presence and absence of workers
attach(donor_effect)
t_overall<-t.test(q1post,q2post, mu=0, alternative="t", paired=T)
t_overall
# t-test for the difference between average egg laying rates between Treatment and Control queens

lm_donor_effect<-lm((q1post - q2post) ~ q3pre * Relatedness, data = donor_effect)             
summary(lm_donor_effect)
# testing for the effects of donor queen fecundity and relatedness between donor and treatment queens on the difference in fecundity between treatment and control queens
hist(resid(lm_donor_effect))
shapiro.test(resid(lm_donor_effect)) 
# check distribution of residuals
interact_plot(lm_donor_effect, pred = q3pre, modx = Relatedness, plot.points=T, interval=T, int.width=0.95,  x.label = "Relatedness between Treatment Queen and Donor Queen",y.label = "Treatment Queen fecundity - Control queen fecundity", modx.values = c(0.36,0.63,0.9))+ theme_classic()
# Plotting Figure 1


cannibalism<-read.table("canni.txt", header=T)
 # read data file with cannibalism observations
names(cannibalism)
 # four variables: colony identity, queen treatment, and numbers of eggs alive and presumed dead
attach(cannibalism)
Model1<- glmer(cbind(Dead, Alive) ~ Queen + (1|as.factor(colony)), data=cannibalism, family = binomial)
# model with treatment as an explanatory variable
Model2<- glmer(cbind(Dead, Alive) ~ 1 + (1|as.factor(colony)), data=cannibalism, family = binomial)
# model without treatment
anova(Model1,Model2)
# ANOVA testing for the effect of treatment
summary(glht(Model1,mcp(Queen="Tukey")))
# pairwise comparisons using Tukey's test

for_plot <- ddply(cannibalism, "Queen", summarise,
               N    = length(Alive),
               mean = mean(Alive),
               sd   = sd(Alive),
               se   = sd / sqrt(N),
               ci = qt ((1-0.05)/2+0.5, N-1) * se
)
ggplot(for_plot, aes(x = Queen, y = mean)) +
  geom_bar(stat = "identity",color="black",fill="white")  +
  theme_bw()+
  theme(panel.grid.major = element_blank(), panel.grid.minor = element_blank())+
  geom_errorbar(aes(ymin = mean - se, ymax = mean + se), width = 0.2)+
  xlab("Queen treatment")+
  ylab("Number of surviving eggs (mean & S.E.M)")
#plotting Figure 2
